# Therapeutic targeting of ACLY in T-ALL *in vivo*

**DOI:** 10.1101/2023.03.27.534395

**Authors:** Victoria da Silva-Diz, Amartya Singh, Maya Aleksandrova, Oekyung Kim, Christopher Thai, Olga Lancho, Patricia Renck Nunes, Hayley Affronti, Xiaoyang Su, Kathryn E. Wellen, Daniel Herranz

## Abstract

T-cell Acute Lymphoblastic Leukemia (T-ALL) is a hematological malignancy in need of novel therapeutic approaches. Here, we identify the ATP-citrate lyase ACLY as overexpressed and as a novel therapeutic target in T-ALL. To test the effects of ACLY in leukemia progression, we developed an isogenic model of NOTCH1-induced *Acly* conditional knockout leukemia. Importantly, we observed intrinsic antileukemic effects upon loss of ACLY, which further synergized with NOTCH1 inhibition *in vivo.* Metabolomic profiling upon ACLY loss revealed a metabolic crisis with reduced acetyl-CoA levels, as well as a decreased oxygen consumption rate. Gene expression profiling analyses showed that the transcriptional signature of ACLY loss very significantly correlates with the signature of MYC loss *in vivo*. Mechanistically, the decrease in acetyl-CoA led to reduced H3K27ac levels in *Myc*, resulting in transcriptional downregulation of *Myc* and drastically reduced MYC protein levels. Interestingly, our analyses also revealed a reciprocal relationship whereby *ACLY* itself is a direct transcriptional target of MYC, thus establishing a feedforward loop that is important for leukemia progression. Overall, our results identified a relevant ACLY-MYC axis and unveiled ACLY as a novel promising target for T-ALL treatment.

## Introduction

T-cell acute lymphoblastic leukemia (T-ALL) is an aggressive hematological disease of immature T-cell progenitor cells. Although most patients nowadays get cured thanks to intensive chemotherapy regimens, 20-50% of patients still relapse and have dismal prognosis^1^. Targeting different metabolic routes in patients has resulted in notable antileukemic effects, as exemplified by the clinical use of methotrexate, 6-mercaptopurine or L-asparaginase^2^. Moreover, we recently identified the antileukemic effects of either serine hydroxymethyltransferase inhibition^3^ or mitochondrial uncoupling^4^, highlighting the value of further exploring metabolic vulnerabilities in leukemia. We previously demonstrated that the metabolic consequences of NOTCH1 inhibition in T-ALL are critical for its antileukemic effects^5^. Since NOTCH1 activating mutations occur in over 60% of patients^6^, we hypothesized that thorough analyses of the metabolic enzymes being downregulated upon NOTCH1 inhibition might yield novel therapeutic targets for T-ALL treatment and identified the ATP citrate-lyase *ACLY*, a critical metabolic and epigenetic regulator^7^, as significantly downregulated upon NOTCH1 inhibition, suggesting a relevant role for ACLY in NOTCH1-driven leukemia. However, nothing is known about the role of ACLY in T-cell development or T-ALL.

## Methods

### Mice

Generation of NOTCH1-driven conditional inducible knockout leukemias was done as previously described^5^. All animal housing, handling, and procedures involving mice were approved by the Rutgers Institutional Animal Care and Use Committee, in accordance with all relevant ethical regulations.

### Profiling experiments

Transcriptional, epigenetic, and metabolomic profiling experiments were done as previously described^8^.

### Molecular biology experiments

Western Blot and luciferase reporter assays were done following standard procedures, as previously described^8^. Detailed methods are described in Supplemental Methods.

## Results and discussion

### ACLY is overexpressed in T-ALL

We previously demonstrated the critical effects of the metabolic consequences of NOTCH1 inhibition *in vivo* for its antileukemic effects^5^. Interestingly, we found the ATP-citrate lyase *ACLY* among the metabolic genes downregulated by NOTCH1 inhibition in T-ALL *in vivo* (**Figure 1A**). ACLY has been previously suggested as an important therapeutic target in solid tumors such as pancreatic cancer^9^, and recent studies have described certain roles for ACLY in the normal physiological function of CD4 and CD8 T-cells^10,11^. However, its putative role in the normal development of T-cells and other hematological lineages, as well as in leukemic transformation and progression, remains largely unknown. ACLY controls both metabolic and epigenetic processes, given its critical role as the rate-limiting enzyme in fatty acid synthesis, as well as by its generation of acetyl-CoA (critical for histone acetylation) as a product of its catalytic reaction^7^. Since both metabolic and epigenetic mechanisms of resistance to NOTCH1 inhibition in T-ALL have been previously described^5,8,12^, our findings suggested a potential role for ACLY in the response to NOTCH1 inhibition and, more broadly, as a relevant player in T-ALL. Next, we analyzed available gene expression data from T-ALL patients^13^ and observed a broad upregulation of *ACLY* as compared to normal T-cell subsets (**Figure 1B**). In line with this, *ACLY* levels were similar across all different T-ALL clinical subgroups^14^. However, samples with NOTCH1 or FBXW7 mutations showed a trend towards increased *ACLY* levels, which reached statistical significance in the cases with simultaneous NOTCH1/FBXW7 mutations (**Supplemental Figure 1A-B**). ACLY overexpression in T-ALL was confirmed at the protein level, with human T-ALL cell lines showing higher expression than normal human thymus or peripheral blood cells (**Figure 1C**). Taken together, these results suggest a previously unknown but important role for ACLY in T-ALL.

**Figure 1.**
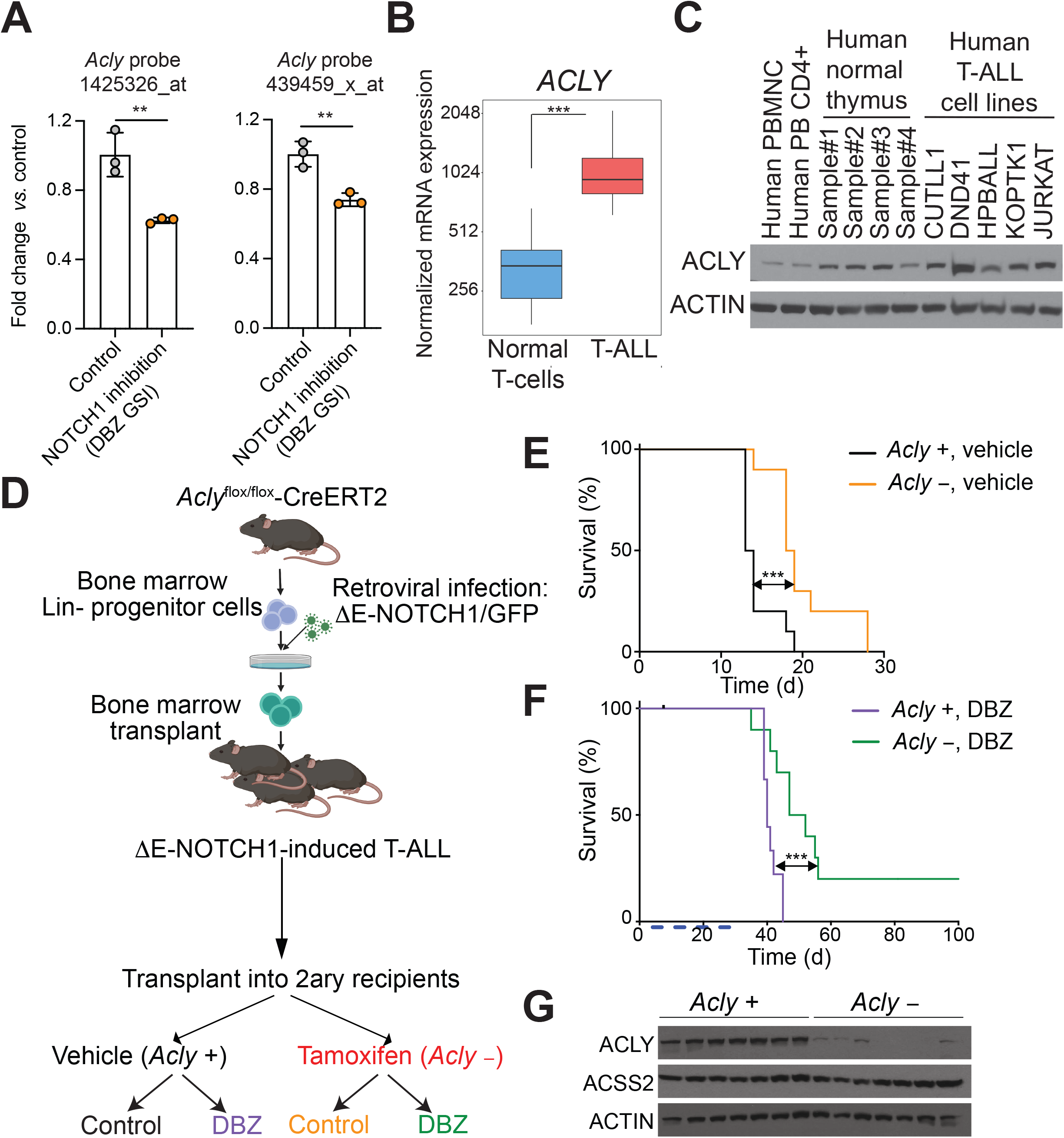
ACLY is a novel therapeutic target in T-ALL. (A) mRNA levels of *Acly* upon NOTCH1 inhibition with the DBZ gamma-secretase inhibitor in mouse T-ALL *in vivo*^5^ (\*\**P*<0.01 calculated with 2-tailed Student’s *t*-test). (B) Box-plot showing *ACLY* expression among T-ALL samples (n=57) and physiological thymocyte subsets (n=21)^13^. Quantile normalization was performed across samples. Boxes represent first and third quartiles and lines represent the median. Whiskers represent the upper and lower limits (*P*<0.001 using Mann-Whitney U-Test; FDR<0.05 using Benjamini-Hochberg correction). (C) Western blot analysis of ACLY and ACTIN expression in human peripheral blood mononuclear cells (PBMNC), CD4+ T-cells, or healthy human thymocytes, as compared to human T-ALL. (D) Schematic illustration of retroviral-transduction protocol for the generation of NOTCH1-induced T-ALLs from inducible *Acly*-conditional knockout mice, followed by transplant into secondary recipients treated with vehicle (*Acly*^+/+^) or tamoxifen (*Acly*^−/−^), with or without DBZ. (E) Kaplan-Meier survival curves of mice harboring *Acly*-positive and *Acly*-deleted isogenic leukemias (n = 10 per group). (F) Kaplan-Meier survival curves of mice harboring *Acly*-positive and *Acly*-deleted isogenic leukemias treated with 4 cycles DBZ (5 mg/kg) on a 4-days-ON (blue blocks at the bottom) and 3-days-OFF schedule (n = 10 per group). (G) Western blot analysis of ACLY, ACSS2, and ACTIN expression in leukemic spleens from terminally ill mice from the survival curve in Figure 1E. \*\*\**P* < 0.005 in panels B and C calculated with log-rank test; \**P* < 0.05; \*\**P* < 0.01; \*\*\**P* < 0.005 in panels D-F calculated with log-rank test calculated with two-tailed Student’s *t*-test.

### ACLY is critical for T-ALL progression but dispensable for T-cell development

Next, we hypothesized that targeting ACLY in T-ALL might confer therapeutic effects. To test this hypothesis, we generated a model of NOTCH1-induced *Acly* conditional knockout leukemia using a well-established protocol^5^ of retroviral transduction of an oncogenic form of NOTCH1 in bone marrow progenitor cells from *Acly* conditional knockout mice, followed by transplantation into lethally irradiated recipients (**Figure 1D**). In time, these mice developed NOTCH1-induced leukemias, which were subsequently transplanted into a secondary cohort of mice and treated with vehicle control or tamoxifen, to induce isogenic loss of *Acly*. Mice were further subdivided into control-treated or treated with the gamma-secretase inhibitor DBZ, to test the response to NOTCH1 inhibition *in vivo* with or without ACLY expression (**Figure 1D**). Notably, our results demonstrated intrinsic antileukemic effects for ACLY loss (**Figure 1E**), which further synergized with NOTCH1 inhibition resulting in a 20% cure rate of leukemic mice (**Figure 1F**). Interestingly, leukemias eventually developing in tamoxifen-treated mice were not genetic escapers and did not show a concomitant feedback upregulation of ACSS2 (**Figure 1G**). ACSS2 upregulation has been previously shown to mediate resistance to ACLY loss in certain contexts^15^, suggesting that T-ALL cells can still survive ACLY loss via alternative mechanisms.

Many of the relevant therapeutic targets of T-ALL progression also show an effect in normal T-cell development, and the potential effects of ACLY loss in normal healthy hematopoietic cells would also be important to predict potential toxicities associated to therapies aimed at inhibiting ACLY. To this end, we bred the *Acly* conditional knockout mice with Vav-iCre expressing mice. Mice with hematopoietic-specific ACLY loss were viable and showed no overt phenotype. Detailed immunophenotypic analyses of thymi from these mice showed no differences in thymus weight or cellularity (**Supplemental Figure 2A-B**), and we also failed to detect any significant difference in the numbers or proportions of different thymocyte subsets (**Supplemental Figure 2C-F**). Overall, our data suggest that ACLY is dispensable for normal T-cell development but critical for T-ALL progression and, thus, may constitute a novel and attractive therapeutic target.

### ACLY loss leads to a metabolic crisis and reduced tumor burden

To explore the mechanistic effects of ACLY loss, we next performed an acute deletion experiment in which mice harboring *Acly* conditional knockout leukemias were first allowed to become fully leukemic before treating them with vehicle (control) or tamoxifen (to induce ACLY loss), followed by euthanasia 72h later (**Figure 2A**). In this setting, acute ACLY loss translated into reduced *Acly* levels (**Figure 2B**) and led to reduced tumor burden (**Figure 2C**). Mice harboring *Acly*-deleted leukemias showed significantly reduced infiltration across tissues, including the spleen or the bone marrow and, most notably, also showed reduced central nervous system infiltration in the meninges (**Figure 2D**). These results were mainly driven by cytotoxic effects, as revealed by increased AnnexinV staining upon ACLY loss (**Figure 2E**).

**Figure 2.**
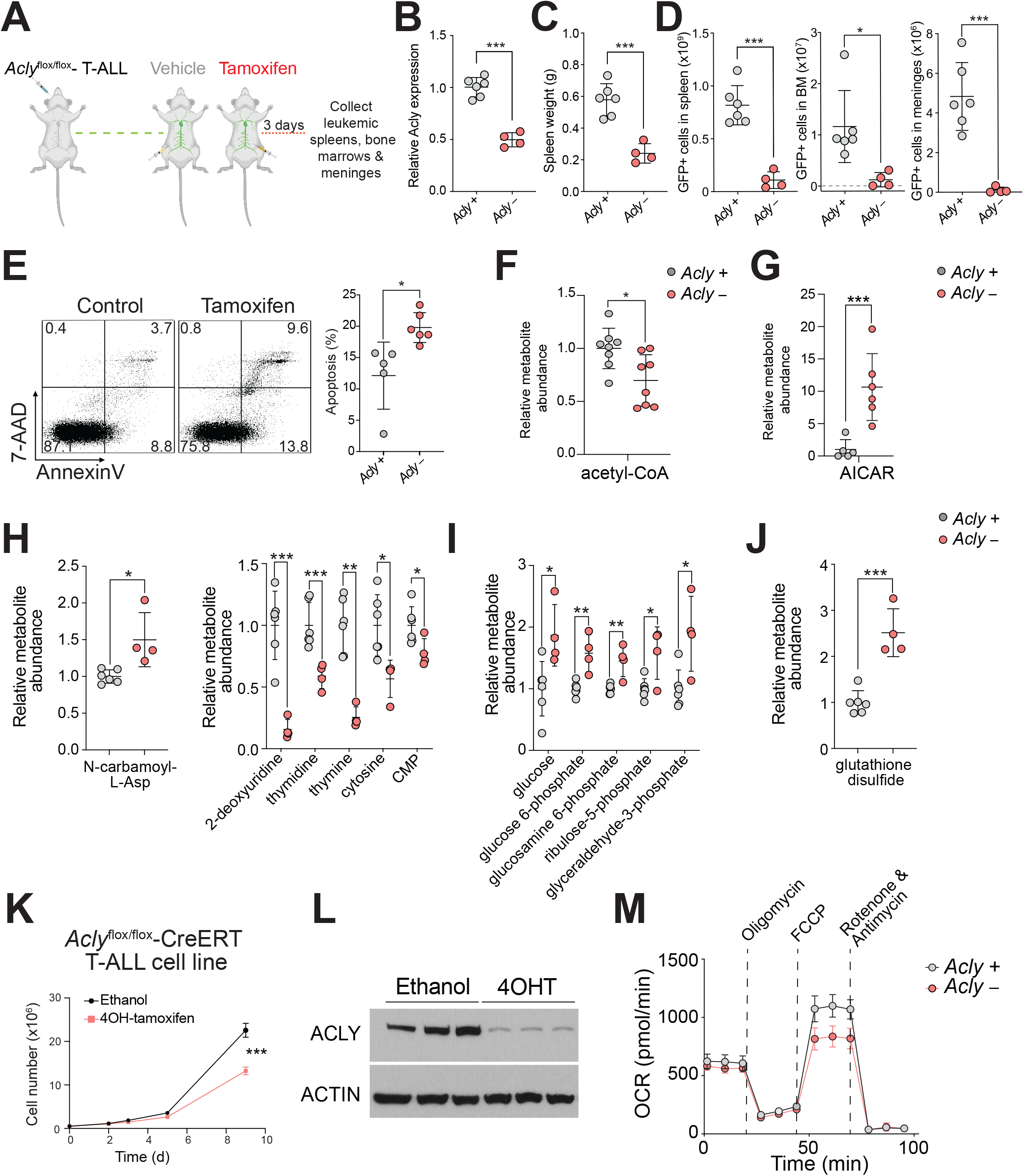
Metabolic consequences of ACLY loss in T-ALL. (A) Schematic of acute ACLY deletion experiment in leukemic mice *in vivo*. (B) Quantitative RT-PCR analysis of *Acly* mRNA expression in tumor cells isolated from ΔE-NOTCH1-induced *Acly* conditional knockout leukemia–bearing mice 72 h after being treated with vehicle only (*Acly*^+/+^) or tamoxifen (*Acly*^−/−^) *in vivo*. (C-D) Tumor burden in ΔE-NOTCH1-induced *Acly* conditional knockout leukemia–bearing mice 72 h after being treated with vehicle only (*Acly*^+/+^) or tamoxifen (*Acly*^−/−^) *in vivo* as revealed by total spleen weight (C) or by the infiltration of GFP-positive leukemic cells in the spleen, bone marrow or meninges (D). (E) Representative flow cytometry plots from of annexin V (apoptotic cells) and 7-AAD (dead cells) staining (left) and quantification of apoptosis (right) in leukemic spleens from ΔE-NOTCH1-induced *Acly* conditional knockout leukemia–bearing mice 48 h after being treated with vehicle only (*Acly*^+/+^) or tamoxifen (*Acly*^−/−^) *in vivo*. (n = 4-5 per treatment in panels B-E; \**P* < 0.05 and \*\*\**P* < 0.005 using two-tailed Student’s *t*-test.) (F-G) Relative Acetyl-CoA (F) and AICAR (G) abundance upon isogenic loss of *Acly* in leukemic spleens 48-72h after being treated with vehicle only (*Acly*^+/+^) or tamoxifen (*Acly*^−/−^) *in vivo*. (H) Relative abundance of the indicated pyrimidine intermediates upon tamoxifen-induced isogenic loss of *Acly* in leukemic spleens from mice treated as in A. (I) Relative abundance of the indicated glycolytic intermediates upon tamoxifen-induced isogenic loss of *Acly* in leukemic spleens from mice treated as in A. (J) Relative abundance of glutathione disulfide upon tamoxifen-induced isogenic loss of *Acly* in leukemic spleens from mice treated as in A. (K-L) Growth curve (K) and western blot analyses of ACLY and ACTIN expression (L) in an *Acly* conditional knockout T-ALL cell line *in vitro* upon treatment with ethanol (control) or 4-hydroxytamoxifen (4OHT) to induce ACLY loss. (M) Oxygen consumption rate (OCR) in *Acly* conditional knockout T-ALL cells, under basal conditions or after 4OHT-induced loss of ACLY, measured in real-time using a Seahorse XF24 instrument. Data are presented as mean ± SD of n = 4 wells. n = 6-8 per treatment in panels F-G and n = 4-5 per treatment in panels H-J; \**P* < 0.05; \*\**P* < 0.01 and \*\*\**P* < 0.005 using two-tailed Student’s *t*-test.

Next, given the well-known metabolic role of ACLY, we performed global metabolomic analyses upon acute ACLY loss *in vivo*. Interestingly, we observed an expected reduction in acetyl-CoA levels (**Figure 2F**), as well as a strong accumulation of AICAR (**Figure 2G**), a purine intermediate and a well-described activator of AMPK^16^. In addition, we also detected accumulation of the pyrimidine intermediate N-carbamoyl-L-aspartate with a concomitant global reduction in pyrimidines (**Figure 2H**), overall suggesting a block in nucleotide biosynthesis. Finally, we also detected an accumulation of glucose and glycolytic intermediates (**Figure 2I**), as well as an increase in oxidized glutathione levels (**Figure 2J**). Consistent with these metabolic effects, loss of ACLY in an *Acly* conditional knockout T-ALL cell line generated from our primary leukemia led to impaired proliferation (**Figure 2K-L**), together with a significant reduction in oxygen consumption rate (**Figure 2M**). Overall, our results suggest that ACLY loss leads to a metabolic crisis in T-ALL cells.

### An ACLY – MYC feedforward loop regulates T-ALL progression

Next, gene expression profiling analyses revealed a strong transcriptional signature upon acute ACLY loss (**Figure 3A** and **Supplemental Table 1**). Pathway analyses using EnrichPlot revealed strong downregulation of pathways related to oxidative phosphorylation and translation (**Figure 3B**), consistent with the metabolic changes previously observed. To gain a deeper understanding of the transcription factor/s that may be governing these transcriptional effects, we analyzed our gene expression profile data using Enrichr^17^ and Dorothea^18^. Intriguingly, both algorithms identified a downregulation of *MYC* activity as potentially related to the changes observed (**Figure 3C** and **Supplemental Figure 3A**). Previous reports suggested a potential regulation of *MYC* by ACLY in endothelial cells^19^, as well as the reverse regulation of *ACLY* by MYC in prostate cancer^20^. To unravel the directionality of the regulation in our T-ALL setting, we performed gene set enrichment analyses (GSEA)^21^ against the signature of MYC loss in T-ALL upon deletion of the N-Me enhancer^22^. Our results showed a very strong correlation between both signatures (**Figure 3D**), and western blot analyses confirmed a drastic downregulation in MYC protein levels (**Figure 3E**), which is consistent with the strong antileukemic effects observed. Given the well-described role of ACLY in epigenetic regulation and our own findings of reduced acetyl-coA levels upon ACLY loss (**Figure 2F**), we next performed ChIP-seq epigenetic profiling experiments, which revealed a global reduction in H3K27ac mark with a concomitant milder reduction in H3K4me3 levels (**Figure 3F**). Interestingly, the effect in histone acetylation might be specific for H3K27ac, as we did not observe major changes in H3K9ac (**Figure 3F**). Importantly, these epigenetic changes were evident in the *Myc* promoter, which showed markedly reduced levels of H3K27ac and a milder decrease in H3K4me3 (**Figure 3G**), consistent with the changes in MYC activity previously observed.

**Figure 3.**
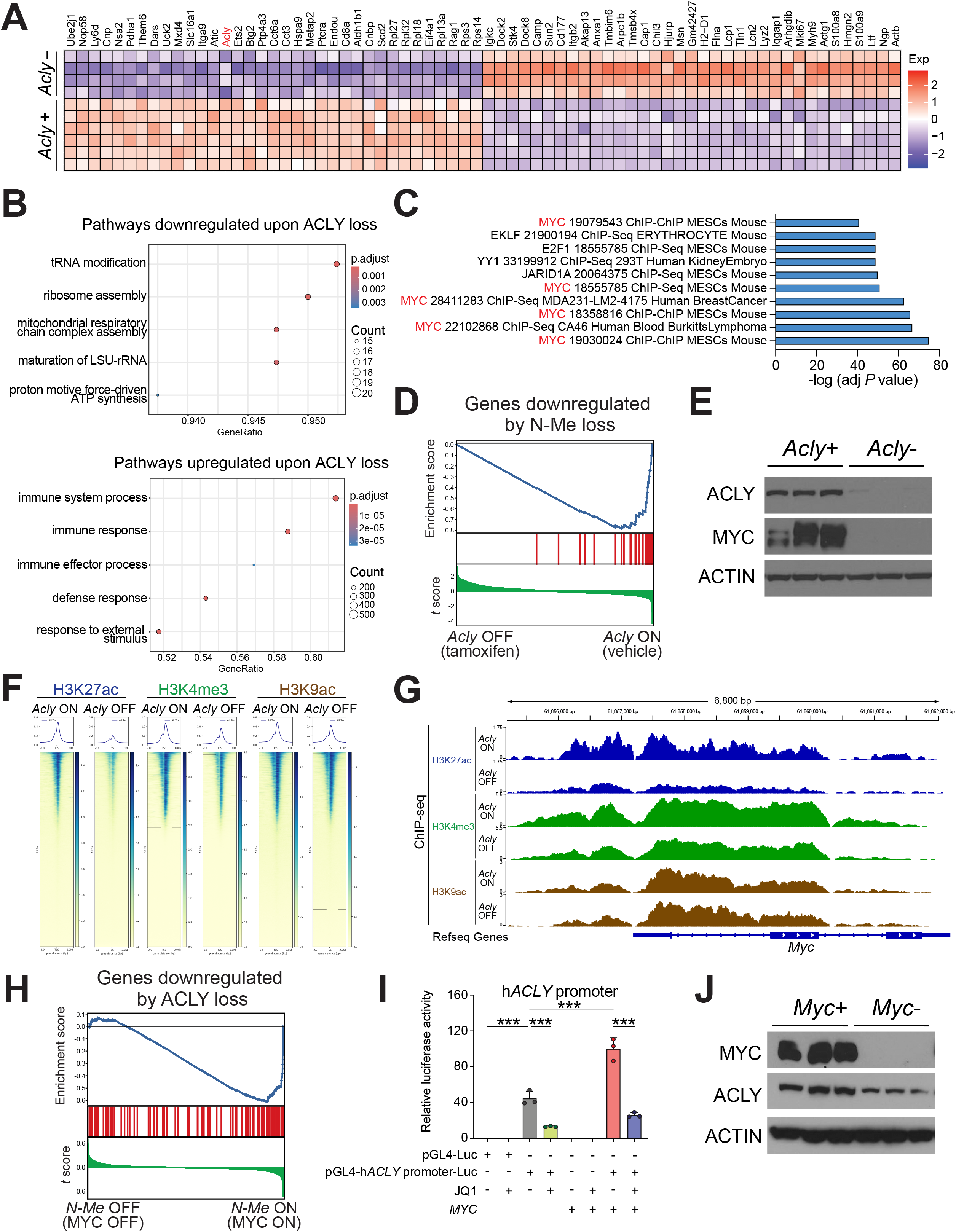
ACLY-MYC axis in T-ALL (A) Heatmap representation of the top differentially expressed genes between control (*Acly*^+/+^) and tamoxifen-treated (*Acly*^−/−^) *Acly*-conditional knockout ΔE-NOTCH1–induced leukemias. Cutoffs used: top 40 up and downregulated genes; *P*-adjusted value < 0.0001; log fold change >1. Scale bar shows color-coded differential expression, with red indicating higher levels of expression and blue indicating lower levels of expression. (B) Pathway analyses of significantly downregulated or upregulated genes upon ACLY loss using EnrichPlot. (C) Enrichr analyses of transcription factors controlling downregulated pathways upon ACLY loss. (D) GSEA of genes downregulated by N-Me loss in control (vehicle only–treated; ACLY ON) or *Acly*-deleted leukemias (tamoxifen–treated; ACLY OFF) *in vivo.* (E) Western blot analyses of ACLY, MYC, and ACTIN expression 72 h after tamoxifen-induced deletion of *Acly in vivo.* (F) Heatmap representation of genome-wide enrichment for H3K27ac (left), H3K4me3 (middle), and H3K9ac (right) marks in control (vehicle only–treated; ACLY ON) or *Acly*-deleted leukemias (tamoxifen–treated; ACLY OFF) *in vivo.* The color of the heat map indicates the number of CPM-normalized reads within +/-3000bp of annotated Transcription Start Site (TSS) for each gene. (G) Epigenetic profiling around the *Myc* promoter showing ChIP-seq tracks for the indicated histones marks in mouse T-ALL leukemic samples treated as in F. (H) GSEA of genes downregulated by ACLY loss in control (vehicle only–treated; N-Me ON) or N-Me deleted leukemias (tamoxifen–treated; N-Me OFF) *in vivo*^22^. (I) Luciferase reporter activity in 293T cells of a pGL4 promoter empty construct (pGL4-Luc) or a pGL4 promoter plus the human *ACLY* promoter, in the presence or absence of MYC overexpression or treatment with JQ1. Data from three independent transfection replicates are shown. \*\*\**P* < 0.005 using two-tailed Student’s *t*-test. (J) Western blot analyses of ACLY, MYC, and ACTIN expression 48 h after tamoxifen-induced deletion *in vitro* of N-Me using an N-Me conditional knockout T-ALL cell line^22^.

Conversely, reverse GSEA analyses also showed that the signature of MYC loss upon N-Me deletion is very similar to the one obtained upon ACLY loss (**Figure 3H**), suggesting that MYC might also be upstream of *Acly*. Indeed, luciferase reporter assays testing this regulation showed that MYC overexpression positively regulates the transcriptional activity of the *ACLY* promoter, whereas treatment with the BRD4 inhibitor JQ1, which prominently inhibits MYC activity^23^, resulted in reduced transactivation of *ACLY* (**Figure 3I**). In line with this, epigenetic profiling of the *ACLY* promoter using publicly available ChIP-seq data in human T-ALL cells uncovered the binding of multiple relevant T-cell-specific transcription factors, including both MYC and NOTCH1 (**Supplemental Figure 3B**). Finally, we took advantage of our mouse N-Me conditional knockout primary T-ALL cell line^22^ to demonstrate that MYC loss in T-ALL resulted in significantly reduced ACLY protein levels (**Figure 3J**), further supporting that MYC also regulates *Acly*. This is consistent with the broad ACLY overexpression observed in T-ALL and with its downregulation upon NOTCH1 inhibition, given the tight link between NOTCH1 and MYC, and the critical role of MYC in T-ALL^22^.

### Resistance to ACLY loss is associated with restored MYC levels

Even if targeting ACLY shows strong intrinsic antileukemic effects *in vivo*, tamoxifen-treated leukemic mice eventually relapse and succumb, and these leukemias are not genetic escapers nor show compensatory upregulation of ACSS2 (**Figure 1**), supporting alternative mechanisms of resistance. Interestingly, western blot analyses of relapsing leukemias showed restored expression of MYC protein levels (**Supplemental Figure 3C**). Similarly, gene expression profiling analyses of relapsing leukemias confirmed the restoration of MYC-driven transcriptional programs and additional pathways that were significantly changing upon acute ACLY loss (**Supplemental Figure 3D-E**).

Collectively, our results identify a therapeutically relevant ACLY-MYC feedforward loop in T-ALL (**Supplemental Figure 4**) in which ACLY provides acetyl-coA units needed for the sustained transcriptional regulation of *MYC*, whereas MYC itself further reinforces this loop by transcriptionally upregulating *ACLY* (**Supplemental Figure 4**). Importantly, while our results identified a critical role for ACLY in T-ALL progression *in vivo*, ACLY is dispensable for normal T-cell development, and mice lacking ACLY in the heme compartment show no gross abnormalities, suggesting that targeting ACLY might have limited toxicities for normal healthy cells. MYC and AKT are two of the most relevant oncogenic factors downstream of NOTCH1 in T-ALL^5,22^. Intriguingly, AKT has also been previously shown to directly phosphorylate and positively regulate ACLY^24^. Thus, the transcriptional program controlled by NOTCH1 in T-ALL leads to increased ACLY activity via multiple pathways, highlighting the key role of ACLY in leukemogenesis. Overall, our results demonstrate that targeting ACLY constitutes a novel and attractive therapeutic option for leukemia treatment. While several ACLY inhibitors have been described to date, these are either relatively weak and unspecific inhibitors, such as bempedoic acid or BMS-303141, or potent but not cell-permeable inhibitors, such as NDI-091143^25^. In this context, our unequivocal genetic results demonstrating a therapeutic role for targeting ACLY in leukemia *in vivo* will hopefully spur additional efforts to develop clinical-grade ACLY inhibitors that can be used in leukemia patients in the near future.

## Acknowledgements

We are grateful to Adolfo A. Ferrando (Columbia University Medical Center) for sharing with us normal human thymocyte samples, and we thank Adolfo A. Ferrando, Antonio Maraver (IRCM, Montpellier), and Laura Belver (IJC, Barcelona) for their constant constructive criticism and support. We also thank everyone involved with JuanLord for their support. Figures 1D and 2A were created using BioRender.com.

## Author contributions

V.dS. performed most molecular biology experiments. A.S. and C.T. performed all computational analyses. M.A., O.K., O.L., P.R.N., and H.A. assisted with several experiments. X.S. supervised metabolomic analyses. K.E.W. contributed critical reagents and gave critical input. D.H. designed the study, supervised the research, and wrote the manuscript together with V.dS., with input from all authors.

