## Supplemental Figures and Methods for "Therapeutic targeting of ACLY in T-ALL *in vivo*"

### *In vivo models of NOTCH1-driven mouse T-ALLs*

Animals were maintained in ventilated caging in specific pathogen-free facilities at New Brunswick RBHS Rutgers Campus. All animal housing, handling, and procedures involving mice were approved by Rutgers Institutional Animal Care and Use Committee (IACUC), in accordance with all relevant ethical regulations.

In order to generate *Acly* conditional inducible knockout leukemias, we used bone marrow progenitor cells (fresh cells provided by K.E.W.) from *Acly*$^{flox/flox}$ mice (JAX #043555) harboring a tamoxifen-inducible Cre recombinase from the ubiquitous UBC locus (JAX #007001). Then, we performed retroviral transduction of lineage-negative enriched cells with a retrovirus encoding a ∆E-NOTCH1-GFP oncogenic activated form of NOTCH1 with concomitant expression of GFP, as previously described[1]. Cells were then transplanted via retro-orbital injection into lethally irradiated (7.5 Gy) recipient mice.

For leukemia-progression studies, already generated ∆E-NOTCH1-GFP-induced *Acly*$^{flox/flox}$-Cre-ERT2/+ leukemias (1 x $10^6$ leukemia cells) were transplanted from primary recipients into sub-lethally irradiated (4.5 Gy) 6-8-week-old secondary recipient C57BL/6 mice (Taconic Farms) by retro-orbital injection. 48 hours after leukemic cell transplantation, recipient mice were treated with vehicle only (corn oil; Sigma, C8267) or tamoxifen (Sigma, T5648; 3 mg per mouse in corn oil), to induce isogenic loss of *Acly* in established leukemias. Subsequently, 5 days post-transplantation, mice in each arm were divided randomly into two different groups: control groups were subsequently treated with vehicle only (2.3 % DMSO in 0.005% methylcellulose, 0.1 % Tween-80) or with DBZ (5 mg per Kg in vehicle solution; Syncom, 29762) on a 4-day-ON and 3-day-OFF schedule, as previously described[1]. Investigators were not blinded to group allocation. Animals were monitored for signs of distress or motor function at least twice daily until they were terminally ill, whereupon they were euthanized.

For *Acly* acute deletion analyses in mouse primary leukemias, we transplanted lymphoblasts from spleens of *Acly* conditional knockout NOTCH1-induced T-ALL-bearing mice into a secondary cohort of recipient mice, as before. We monitored mice until they presented clear leukemic signs with >60% GFP-positive leukemic cells in peripheral blood; then, mice were treated with vehicle or tamoxifen and, 48 or 72 hours after treatment, mice were euthanized, and spleen samples were collected for further analyses.

For T-cell development, we bred *Acly* conditional knockout mice with Vav-iCre mice (JAX #008610) to obtain the different genotypes of interest, and T-cell development was analyzed in 6–8-week-old mice.

***Western blotting***

Human peripheral blood mononuclear cells (PBMCs) and peripheral blood CD4-positive cells were purchased from Lonza (product numbers CC-2702 and 2W-200, respectively). Samples from normal human thymus were kindly provided by Dr. Adolfo Ferrando (Columbia University). Whole-cell extracts were prepared using standard procedures. After protein transfer, membranes were incubated with the antibodies anti-ACLY (1:1000, 15421-1-AP, Proteintech), anti-ACSS2 (1:1000, 3658, Cell Signaling), anti-c-MYC (1:1000, 13987, Cell Signaling), and anti-β-actin-HRP (1:50000; A3854, Sigma). Antibody binding was detected with a secondary antibody coupled to horseradish peroxidase (Sigma, NA934) using enhanced chemiluminescence (Thermo Scientific, 34578).

***Oxygen Consumption Rate (OCR)***

OCR was assessed using an XF24 Seahorse Biosciences extracellular flux analyzer (Agilent Technologies) following the manufacturer's instructions. Briefly, mouse T-ALL cells were resuspended in Seahorse XF RPMI Medium (Agilent Technologies, 103576-100) supplemented with 10 mmol/L glucose (Agilent Technologies, 1003577-100), 1 mmol/L pyruvate (Agilent Technologies, 1003578-100), and 2 mmol/L glutamine (Agilent Technologies, 1003579-100). A density of $1 \times 10^6$ cells per well was seeded into XF24 Seahorse Biosciences plates pre-coated with Cell-Tak (Corning, 354240) and centrifuged to ensure complete attachment. OCR was evaluated through the sequential addition of 1 μmol/L oligomycin (Sigma, O4876), 1 μmol/L FCCP (Sigma, C2920), and 0.5 μmol/L rotenone (Sigma, R8875) and antimycin A (Sigma, A8674) into each well.

***Luciferase Reporter Assays***

Reporter assays were performed using a pGL4.10 Vector (Promega, E6651) luciferase construct alone or coupled with the human ACLY promoter sequence (hg38; chr17: 41918547-41919547; DNA sequence was synthesized by Genewiz), cloned in the forward orientation, and following our previously described protocol[2]. Constructs were transfected into 293T together with a pMSCV-IRES-mCherry control vector or a pMSCV-*cMYC*-IRES-mCherry, and a plasmid driving the expression of the Renilla luciferase gene (pCMV-Renilla) used as an internal control. 48 h after transfection, cells were treated with JQ1 (500nM; MedChemExpress, HY-13030) for 24 h as indicated. We measured luciferase activity 72 h after electroporation with the Dual-Luciferase Reporter Assay kit (Promega, E1980).

***Flow cytometry analysis***

To analyze leukemic spleen samples, single-cell suspensions were prepared by disrupting spleens through a 70 µm filter. Red cells were lysed by incubation with ammonium-chloride-potassium lysing buffer (155 mM NH4Cl, 12 mM KHCO3, and 0.1 mM EDTA) for 5 minutes on ice. Apoptotic cells in leukemic spleens were quantified with PE-AnnexinV Apoptosis Detection Kit I (BD Pharmingen, 559763).

To analyze thymic populations, single-cell suspensions of total thymocytes were prepared by disrupting the tissues through a 70-µm filter. Red cells were removed by incubation with ammonium-chloride-potassium lysing buffer (155mM NH4Cl, 12mM KHCO3 and 0.1mM EDTA) for 5 minutes on ice. Single cells were stained with anti-mouse fluorochrome-conjugated antibodies: CD4 PE-eFluor 610 (1:400, RM4-5, Thermo Fisher Scientific), CD8a PE (1:200, 53-6.7, BD Pharmingen), CD44 PerCPC-Cy5.5 (1:400, IM7, Thermo Fisher Scientific), and CD25 Alexa Fluor 488 (1:1000, 7D4, Thermo Fisher Scientific). Total thymocytes were visualized in a CD4 versus CD8a plot and CD4 SPs (CD4+CD8a–), CD8 SPs (CD4–CD8a+), CD4/CD8 DPs (CD4+CD8a+) and CD4/CD8 DNs (CD4–CD8a–) were gated. To differentiate specific stages of early T-cell development, CD4/CD8 DNs were plotted in a CD44 versus CD25 plot to distinguish DN1 (CD44+CD25–), DN2 (CD44+CD25+), DN3 (CD44–CD25+) and DN4 (CD44–CD25–) populations. All flow cytometry data were acquired using an Attune NxT Flow Cytometer (ThermoFisher Scientific) and analyzed with FlowJo v10.6.2 software (BD).

***Metabolite extraction and LC-MS-based metabolomics***

Leukemic spleen samples were processed and analyzed as previously described[2,3].

***Quantitative RT-PCR***

Total RNA was extracted from cells using RNeasy Plus Mini Kit (Qiagen), and cDNA was generated with High-Capacity cDNA Reverse Transcription Kit (Applied Biosystems). Quantitative PCR was performed on a QuantStudio 12K Flex Real-Time PCR System (Applied Biosystems) using *Power*SYBR Green PCR Master Mix (Applied Biosystems), and transcript levels were normalized to actin as an internal control. Primers used: *Acly* (Forward: 5′-TTCGTCAAACAGCACTTCC-3′; Reverse: 5′-ATTTGGCTTCTTGGAGGTG-3′) and *Actin* (5′-GGCTGTATTCCCCTCCATCG-3′; 5′-CCAGTTGGTAACAATGCCATGT-3′).

***RNA-seq gene expression profiling***

*Acly* conditional knockout ∆E-NOTCH1-induced T-ALL-bearing mice were treated with vehicle only (corn oil; Sigma, C8267) or tamoxifen (Sigma, T5648; 3 mg per mouse in corn oil) to induce isogenic loss of *Acly* via intraperitoneal injection. 72 hours later, single-cell suspensions of total leukemic splenocytes were prepared by pressing leukemic spleens through a 70 μm filter. We removed red cells in spleen samples by incubation with red blood cell lysis buffer (155 mmol/L NH4Cl, 12 mmol/L KHCO3, and 0.1 mmol/L EDTA) for 5 minutes on ice. RNA was extracted using QIAshredder (QIAGEN, 79656) and RNeasy Mini (QIAGEN, 74106) kits. RNA library preparations and next-generation sequencing were performed using Illumina Next-Seq platform (Illumina). We estimated gene-level raw counts and performed differential expression and/or GSEA analyses as previously described[2].

***RNA-seq analysis of Normal vs T-ALL***

*ACLY* expression was analyzed among T-ALL samples (n=57) and physiological thymocyte subsets (n=21) from published literature[4]. Quantile normalization was performed across samples. Differential expression analysis was performed using the Mann-Whitney U-Test and the resultant p-values were corrected for multiple hypothesis testing using the Benjamini-Hochberg correction. Genes with FDR < 0.005 were shortlisted as differentially expressed.

***ChIP-seq Analysis***

Analyses of genome-wide H3K27ac, H3K4me3, and H3K9ac marks in leukemic cells isolated 72 h after vehicle or tamoxifen treatment in leukemic mice harboring *Acly*-conditional knockout leukemias were performed by Active Motif, following well-established protocols and using ChIP-seq validated antibodies. ChIP-seq reads were trimmed using TrimGalore! (v0.6.7) using the default parameters. Thereafter, quality control was performed using Fastqc (v0.11.9). Reads were then aligned to the mm39 reference genome using BWA (v0.7.17) followed by Samtools (v1.17) to sort the aligned reads. Picard (v3.0.0) was used to mark the duplicates. The aligned reads were then filtered using BAMtools (v2.5.2) to remove reads mapped to blacklisted region, reads marked as duplicates, reads that were not marked as primary alignments, reads that were unmapped, and reads mapped but with a low mapping quality (multiple hits, secondary alignments, etc.). Normalized and Relative Strand Cross-correlation (NSC/RSC) was computed using Phantompeakqualtools (v 1.2.2). BigWig files were then created using deepTools (v 3.5.1). Finally, both narrow and broad peak calling was performed using MACS2 (v 2.7.1) with the q-value threshold set to 0.1. Differential peak analysis was performed using edgeR (v 4.20).

***Statistical analysis***

Statistical analyses were performed with Prism 8.0 (GraphPad). Unless otherwise indicated in figure legends, statistical significance between groups was calculated using an unpaired two-tailed Student's *t*-test. Survival in mouse experiments was represented with Kaplan–Meier curves, and significance was estimated with the log-rank test.

***Data availability***

RNA-seq and ChIPseq data from *Acly* conditional knockout leukemias have been deposited in the NCBI Gene Expression Omnibus (GEO) repository. We analyzed epigenetic profiling and transcription factor binding using the following human T-ALL publicly available datasets from GEO: GSE58406, GSE83777, GSE138516, GSE85524, GSE59657, GSE29600, GSE124223.

**Supplemental Figures and Supplemental Figure Legends**

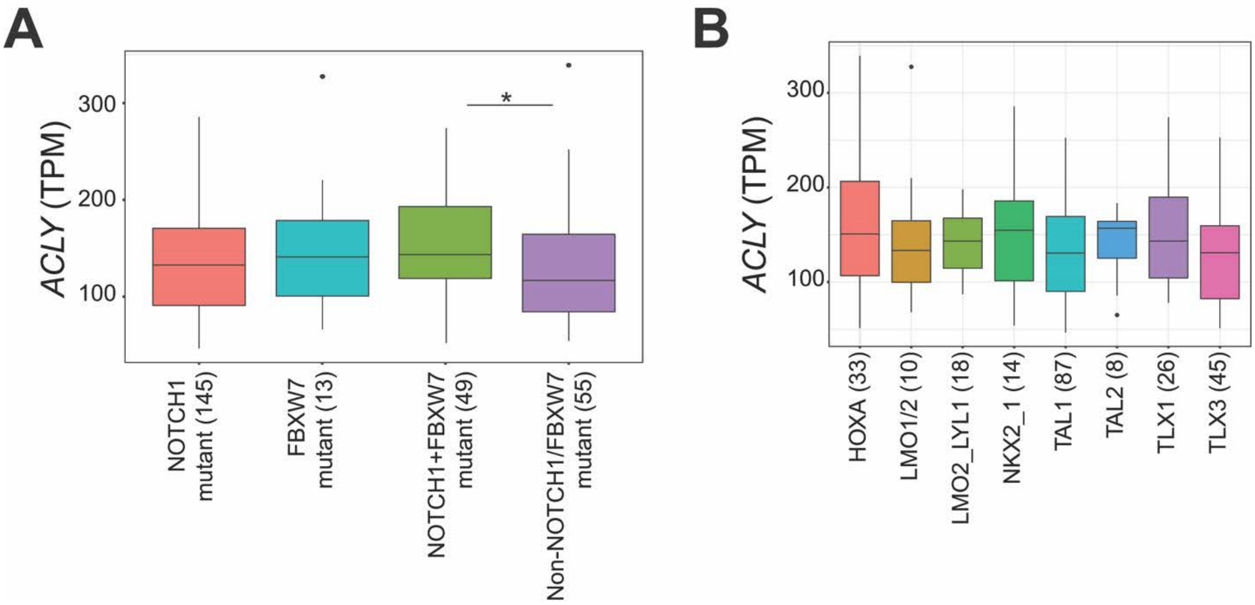


**Supplementary Figure 1**. ACLY is overexpressed in human T-ALL. (A) *ACLY* expression levels in human primary T-ALL cases from different subclinical groups[5]. (B) *ACLY* expression levels in human primary T-ALL cases with/without *NOTCH1* and *FBXW7* mutations[5] (Wilcoxon rank-sum test *$P < 0.05$).

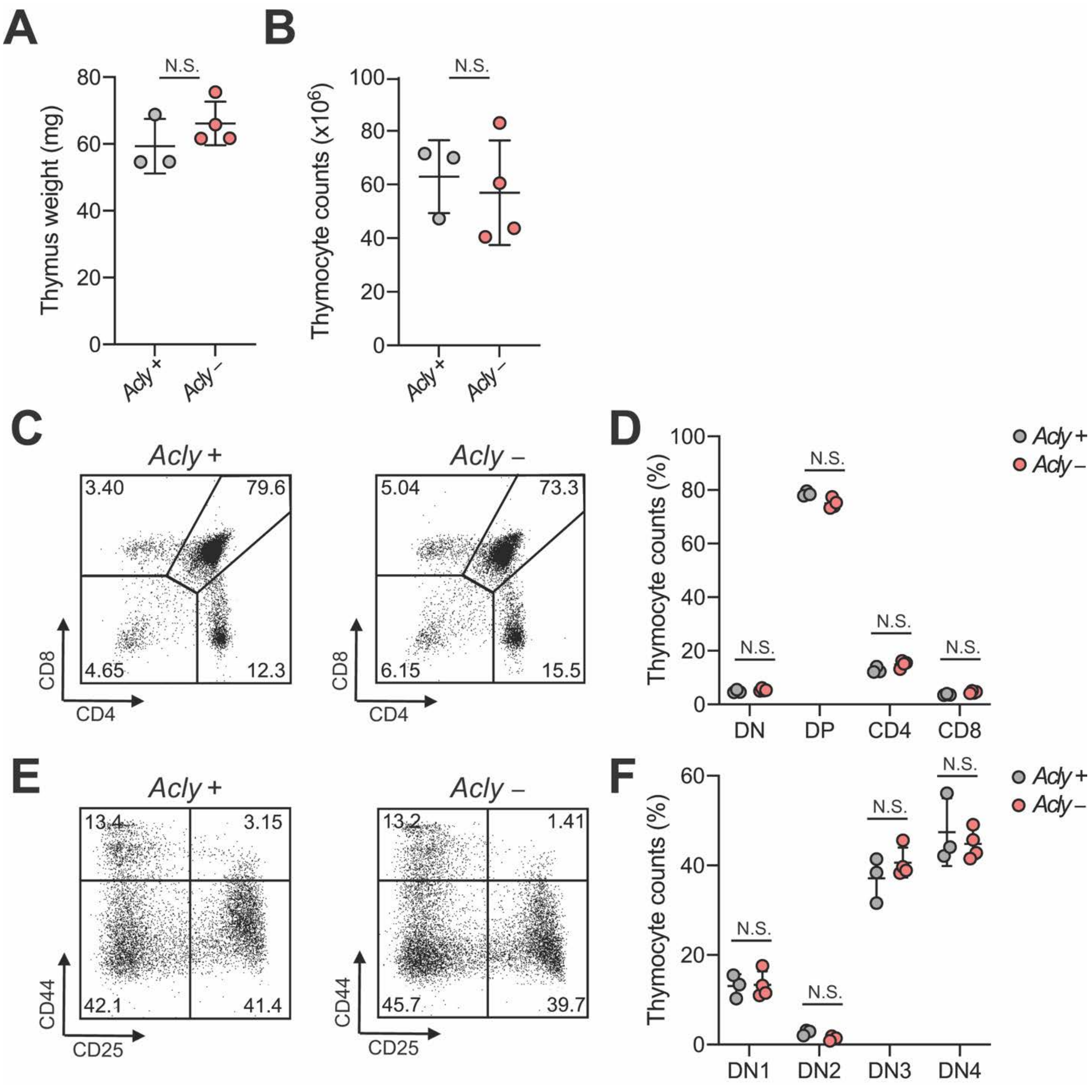


**Supplementary Figure 2.** ACLY loss is dispensable for normal T-cell development *in vivo*. (A) Thymus weight in *Acly*$^{flox/flox}$-Vav-iCre and *Acly*$^{+/+}$-Vav-iCre mice. (B) Total thymocyte count in *Acly*$^{flox/flox}$-Vav-iCre and *Acly*$^{+/+}$-Vav-iCre mice. (C-D) Representative plots (C) and quantification (D) of CD4–CD8– (DN), CD4+CD8+ (DP), CD4+CD8– (CD4) and CD4–CD8+ (CD8) populations in the thymus from *Acly*$^{flox/flox}$-Vav-iCre and *Acly*$^{+/+}$-Vav-iCre mice. (E-F) Representative plots (E) and quantification (F) of

CD44+CD25– (DN1), CD44+CD25+ (DN2), CD44–CD25+ (DN3) and CD25–CD44– (DN4) populations in the thymus from *Acly*$^{flox/flox}$-Vav-iCre and *Acly*$^{+/+}$-Vav-iCre mice. No comparison was significant (N.S.) using two-tailed Student’s *t*-test.

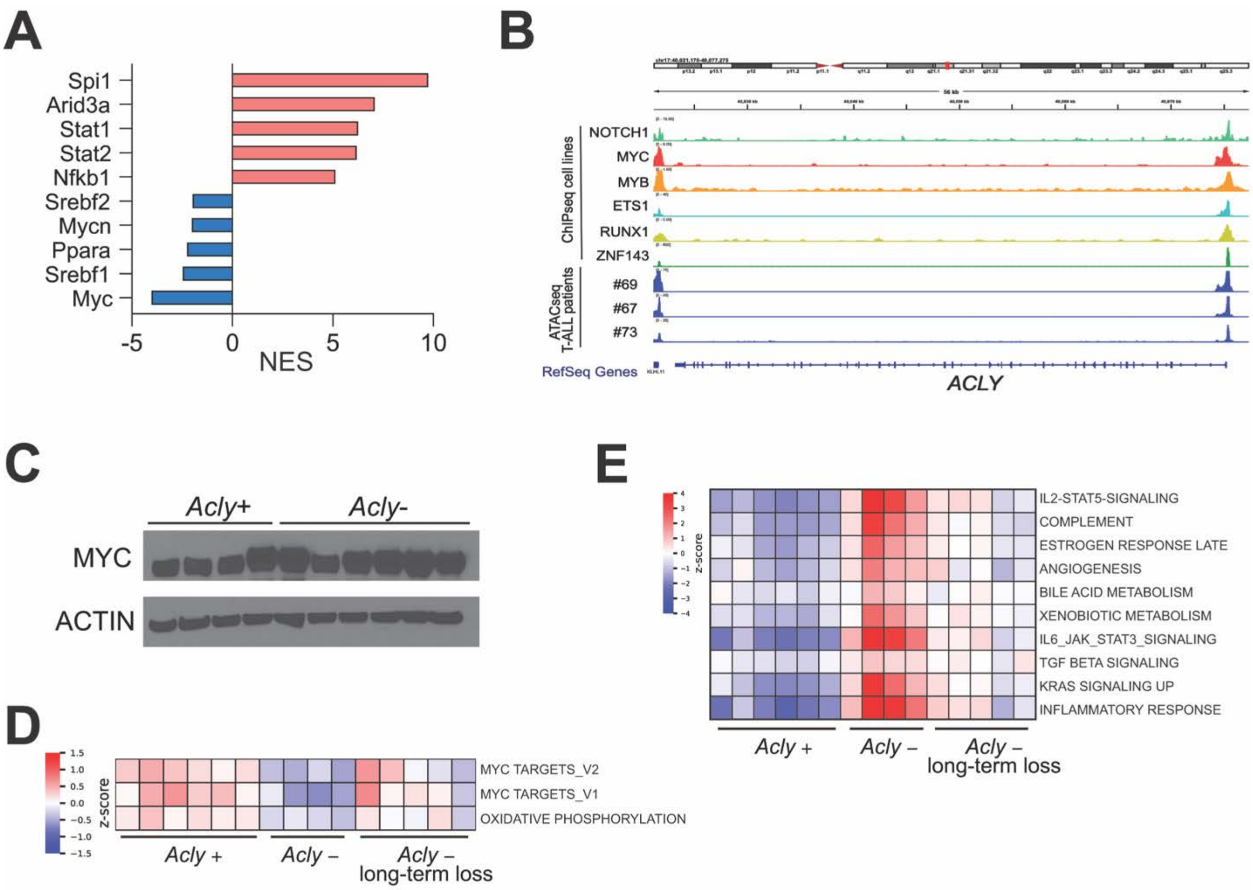


**Supplementary Figure 3.** ACLY-MYC reciprocal axis. (A) Normalized Enrichment Score (NES) analysis using Dorothea algorithm[6] of transcription factors governing the signature of acute ACLY loss in leukemia–bearing mice 72 h after being treated with vehicle only (*Acly* +) or tamoxifen (*Acly* – acute loss) *in vivo*. (B) ChIP-seq and ATAC-seq profiling of the *ACLY* promoter in different human T-ALL cell lines and primary T-ALL patient samples. (C) Western blot analysis of MYC and ACTIN expression in leukemic spleens from terminally ill mice from survival curve in Figure 1E. (D-E) Heatmap representations for the most upregulated (D) and most downregulated (E) Hallmark Gene Sets in mouse T-ALL samples after acute and long-term loss of ACLY using GSEA (nominal $P < 0.2$). Expression is represented by a composite z-score using Stouffer's method.

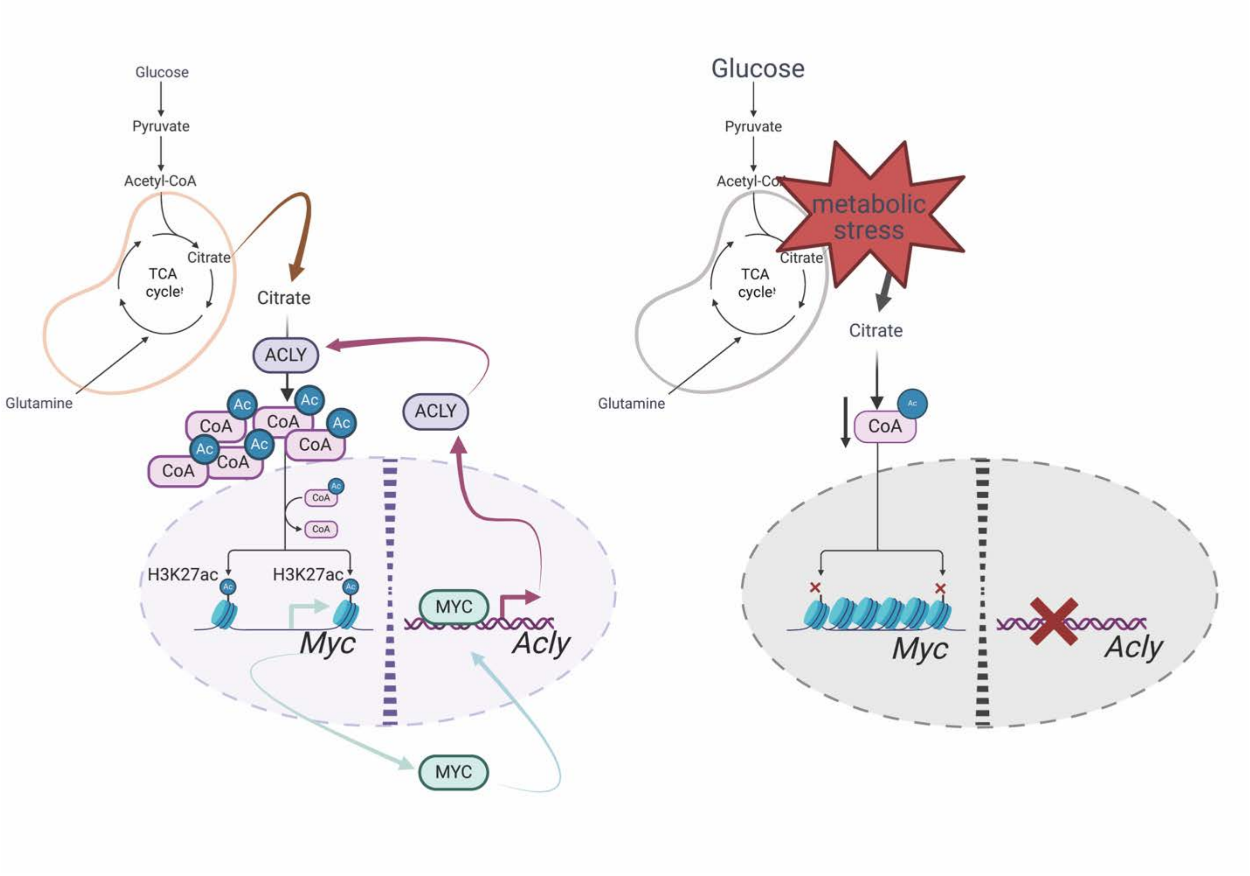


**Supplemental Figure 4.** Schematic representation of the ACLY-MYC axis in T-ALL.
